# Discovery of Large Disjoint Motif in Biological Network using Dynamic Expansion Tree

**DOI:** 10.1101/308254

**Authors:** Sabyasachi Patra, Anjali Mohapatra

## Abstract

Network motifs play an important role in structural analysis of biological networks. Identification of such network motifs leads to many important applications, such as: understanding the modularity and the large-scale structure of biological networks, classification of networks into super-families etc. However, identification of network motifs is challenging as it involved graph isomorphism which is computationally hard problem. Though this problem has been studied extensively in the literature using different computational approaches, we are far from encouraging results. Motivated by the challenges involved in this field we have proposed an efficient and scalable Motif discovery algorithm using a Dynamic Expansion Tree (MDET). In this algorithm embeddings corresponding to child node of expansion tree are obtained from the embeddings of parent node, either by adding a vertex with time complexity O(n) or by adding an edge with time complexity O(1) without involving any isomorphic check. The growth of Dynamic Expansion Tree (DET) depends on availability of patterns in the target network. DET reduces space complexity significantly and the memory limitation of static expansion tree can overcome. The proposed algorithm has been tested on Protein Protein Interaction (PPI) network obtained from MINT database. It is able to identify large motifs faster than most of the existing motif discovery algorithms.

## 1. Introduction

Biological network exhibit both global properties as well as local properties. Some of the global statistical features are small world property, scale-free network characteristics, power law degree distribution etc. Milo et al. [1] first coined the concept of network motif which is treated as local property of biological network. These are statistically over represented patterns having significant functional properties. They constitute the basic building blocks of complex biological networks and essential for functional analysis. Discovery of network motifs is a demanding task in order to define classes of networks and network homologies [1]. Network motif plays a key role in understanding the modularity and the large-scale structure of biological network [2]. They have also been used for networks super family classification [3] and artificial network model for a real-world network, prediction of breast cancer survival outcome [4], cancer disease diagnosis [5], drug repositioning [6] analysis of functional network in diabetes patients [7] etc.

Motifs are recurrent patterns and to estimate its frequency within a network three different frequency measures are widely used: F1, F2 and F3 [8]. Frequency measure F1 allows overlapping of both nodes and edges while counting the matches of a pattern. This concept does not satisfy downward closure property [9]. This indicates that the motif frequency may increase with respect to increase in motif size. Frequency measure F2 allows edge disjoint matches and F3 measure count completely disjoint matches of a pattern. The downward closure property is satisfied by both the frequency measure F2 and F3. Frequency measure F2 is used in this paper as the downward closure property is necessary for pruning the branches of DET.

Network motif is defined as a frequent and unique subgraph pattern in a network [10]. Statistical significance of such unique subgraph pattern is evaluated using p-value and z-score [11]. The z-score is defined as 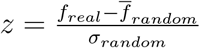; where *f_real_* and 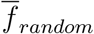 are the frequencies of a motif in the target network and the mean frequency of the motif in randomized networks respectively. *σ_random_* represents the standard deviation of the frequencies in the randomized networks. Higher z-score represents significant motif. The p-value represents the probability that the number of times a motif appears in a randomized network, greater than or equal to the number of times the motif appears in the target network. The lower p-value means significant motif.

Motif discovery has been proved to be a computationally hard problem [12]. Two major challenges of this process are: (1) In order to count frequency of a motif with known topology, it requires to solve subgraph isomorphism problem, which is NP-Complete [13]. (2) Numbers of alternative topology increases exponentially with respect to number of edges in the motif when the motif topology is not known in advance [14]. The existing methods face major challenges when the motif size increases [15–19]. This motivates us to design an efficient and scalable algorithm which can discover large motif in a practical time bound. A motif centric algorithm is proposed that eliminates costly isomorphic test. The central idea of proposed method is to use a Dynamic Expansion Tree (DET) that grows depending on the availability of search pattern in the target network. The expansion tree is initialized with a root node which contains a size-3 tree. Edge disjoint subgraphs corresponding to this root node are computed first. Then the child nodes of the expansion tree are created from the parent node by first adding vertices then edges. Vertex addition continues till the size of the subgraph reaches desired motif size and then edges are added until a complete graph is obtained. Embeddings of the subgraph in the target network are computed along with the growth of expansion tree. A branch of the expansion tree is not expanded further when the frequency of the subgraph failed to cross a threshold. Proposed method developed efficient mechanisms to avoid computationally expensive isomorphism tests during addition of graph elements. Vertex addition can be done with time complexity O(n) and edge addition can be done approximately in time complexity O(1). Evaluation on protein-protein interaction (PPI) networks indicates that proposed method is significantly faster than the existing methods. In addition, memory limitation of static expansion tree is eliminated for large motif discovery.

The rest of this paper is organized as follows: Literature Review is presented in the next section. Proposed method is described next followed by proposed motif discovery algorithm. Then Computational Complexity is discussed. Dataset, experimental evaluation of results and comparison with the existing algorithms are covered in Section Results and discussion. This paper is concluded with a brief conclusion and future direction.

## 2. Literature Review

Some of the research contributions in this domain are discussed below:

The first major contribution in network motif discovery by Milo et al. [1], published in 2002. The significance of the network motif is measured by comparing real network to suitably randomized networks having same degree distribution as real network and found only selected patterns of a real network are statistically significant. A backtracking algorithm, mfinder is used for discovering network motifs. Exponential space complexity of this algorithm made this method incapable to deal with large motif. Kashtan et al. [20] in 2004 improved the execution time of motif finding algorithm by sampling approach, but the results obtained are biased. Sebastian Wernicke [21] in 2005 proposed a specialized algorithm ESU that could avoid redundancy in computation through proper enumeration. This method uses a third-party algorithm NAUTY [22] for checking isomorphism. The flexible pattern finder (FPF) algorithm [8] proposed different frequency concept for computing pattern frequency. The number of patterns grows rapidly with respect to increase pattern size. Therefore, searching all patterns systematically is a time consuming task even for medium-size pattern. The algorithms discussed so far are network centric that discover motifs for the whole network. In 2007, Grochow and Kellis [23] proposed a motif centric algorithm, where frequency counting is done on a specific isomorphic class. This algorithm avoids unnecessary and redundant searches by mapping the query graph only on one representative of its equivalence class. The symmetry conditions are removed by adding constraints on the labelling of the vertices. Kashani et al. [24] in 2009, brought a new network centric algorithm named as Kavosh. It differs from other algorithms in that it builds an implicit tree rooted at the chosen vertex, and then generates all combinations with the desired number of nodes. Omidi et al. proposed MODA [25] in 2009, which is based on a pattern growth methodology. This is a subgraph-centric algorithm. The core idea of this algorithm is to first find the frequency of acyclic subgraphs, save the respective embeddings in memory and then use those embeddings in order to quickly find out the frequencies of cyclic subgraphs. MODA introduces the concept of expansion tree which is static in nature and built at the beginning of the algorithm. A novel algorithm proposed by Cheng Liang et al. [26] in 2015 named as CoMoFinder to accurately and efficiently identify composite network motifs in genome-scale co-regulatory networks. CoMoFinder is developed based on a parallel subgraph enumeration strategy to efficiently and accurately identify composite motifs in large TF-miRNA co-regulatory networks. Rasha Elhesha and Tamer Kahveci [27] in 2016, proposed a motif centric algorithm for finding motifs in a target network. The core idea of this method is to build a set of basic building patterns and find instances of these patterns. Then the size of the motif increased by joining known motifs with the instances of basic building patterns. Lin et al. [28] in 2017 presents a novel study on network motif discovery using Graphical Processing Units (GPUs). The basic idea is to employ GPUs to parallelize a large number of subgraph matching tasks in computing subgraph frequencies from random graphs, so as to reduce the overall computation time of network motif discovery. Yinghan Chen and Yuguo Chen [29] published an efficient sampling algorithm for network motif detection. However sampling approach may lead to biased result. Considering the challenges of existing methods, in the next section a motif centric algorithm is proposed to discover large disjoint network motif.

## 3. Motif Discovery using Dynamic Expansion Tree (MDET)

Pattern growth approach is used in our proposed algorithm. This is a motif centric algorithm where the task is to discover network motifs of a specified size. The central idea of the proposed method is to use a Dynamic Expansion Tree (DET) which regulates motif discovery mechanism. The root node of Expansion Tree (ET) is a minimally connected acyclic graph of three vertices (size-3 tree) and hence the number of embeddings can be computed directly from the adjacency list and adjacency matrix of the target network. The ET grows in two steps; first vertices are added to the basic tree level wise to reach size-k tree. In this step each node of the ET are acyclic graph and the embeddings of these nodes are computed from the embeddings of their parent node using tree census algorithm. In the next step edges are added to the size-k tree in each level till a complete graph is obtained. The embeddings of each node are obtained by graph census algorithm. So before computing the frequency of the query graph present at a particular level of ET, frequency of its parent must be computed and the parent embeddings are obtained from their parent and this process continue in a bottom up manner. Frequency of a node in ET represents the number of embeddings of the subgraph (corresponding to the node) in the target network. In each step edge disjoint embeddings are computed by using MIS algorithm [27]. A node in the ET is expanded only when F2 frequency exceeds the predefined threshold. This pruning criterion is an implication of downward closure property of F2 frequency measure. Hence most of the branches of expansion tree vanishes much before static expansion tree. This feature of DET improves the performance of the algorithm substantially in terms of running time. Before presenting the proposed algorithm we have discussed briefly about static expansion tree, dynamic expansion tree and two key steps used in this algorithm.

### 3.1. Static Expansion Tree (SET)

The concept of expansion tree is used in MODA developed by Omidi et al. [25] in 2009. The Expansion Tree (ET) is represented by *T_k_* where *k* represents the size of target motif. A size-5 dynamic expansion tree is shown in Figure 1.

**Figure 1.**
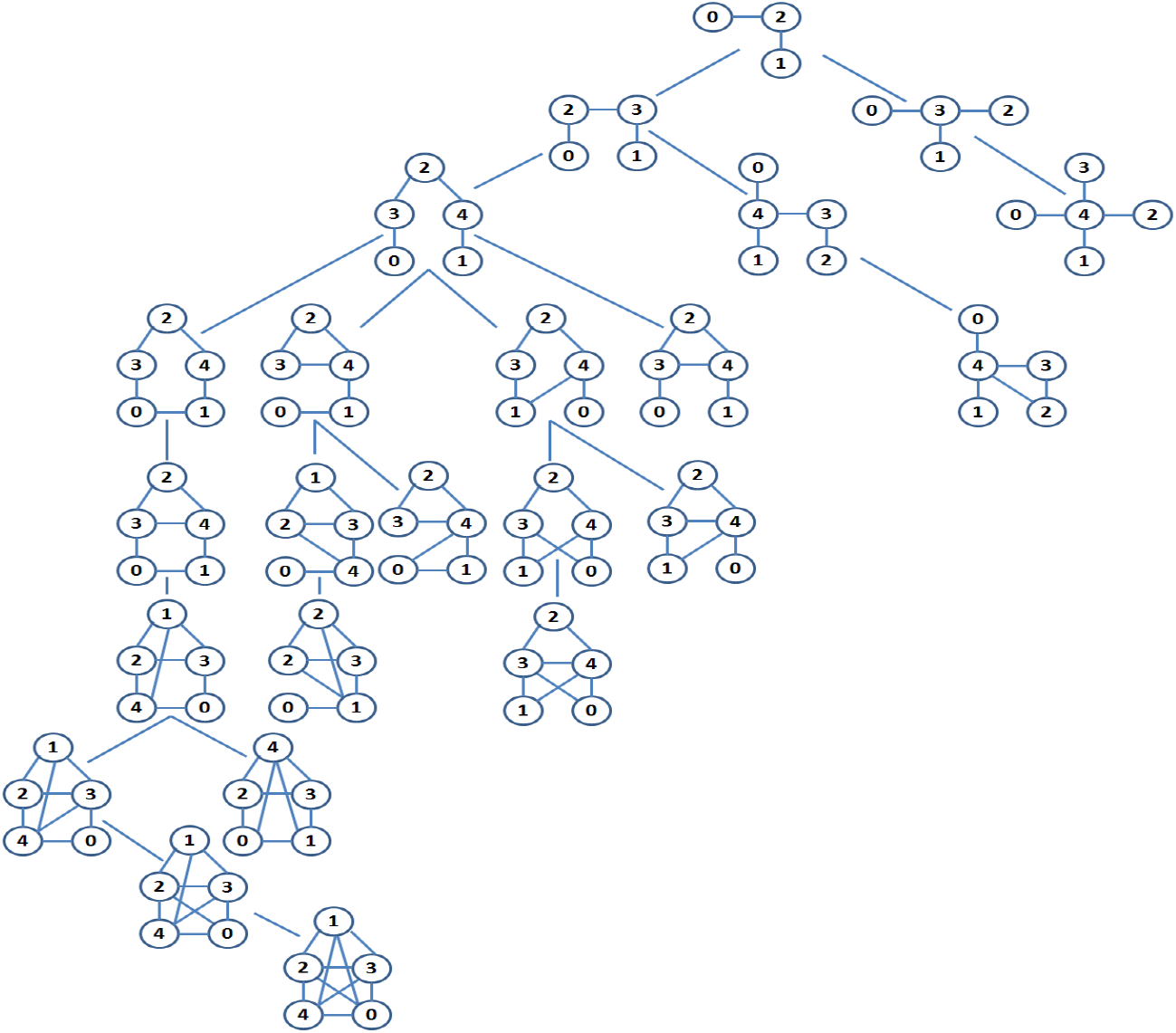
Static expansion tree *T_5_* for size-5 motifs

A size-3 tree is present at level-1. At the second level there are two non-isomorphic size-4 trees, and three non-isomorphic size-5 trees are present at third level. Up to this level, a child graph is obtained by adding a vertex with the parent graph. Isomorphic graph may produce by adding vertices with alternative parent vertex. This is elaborated in detail in vertex addition step. Fourth level onwards an edge is added to the parent graph to form a child graph in each successive level. Similar to vertex addition, alternative edge additions also produce isomorphic graphs. Edge addition continues until a complete graph is obtained.

### 3.2. Dynamic Expansion Tree (DET)

In contrast to static expansion tree, the expansion of Dynamic Expansion Tree (DET) depends on the available motif in the target network. DET also starts with a size-3 tree as root node and grows similar to static expansion tree. However the growth is interrupted by pruning criterion. A size-5 dynamic expansion tree is shown in Figure 2.

**Figure 2.**
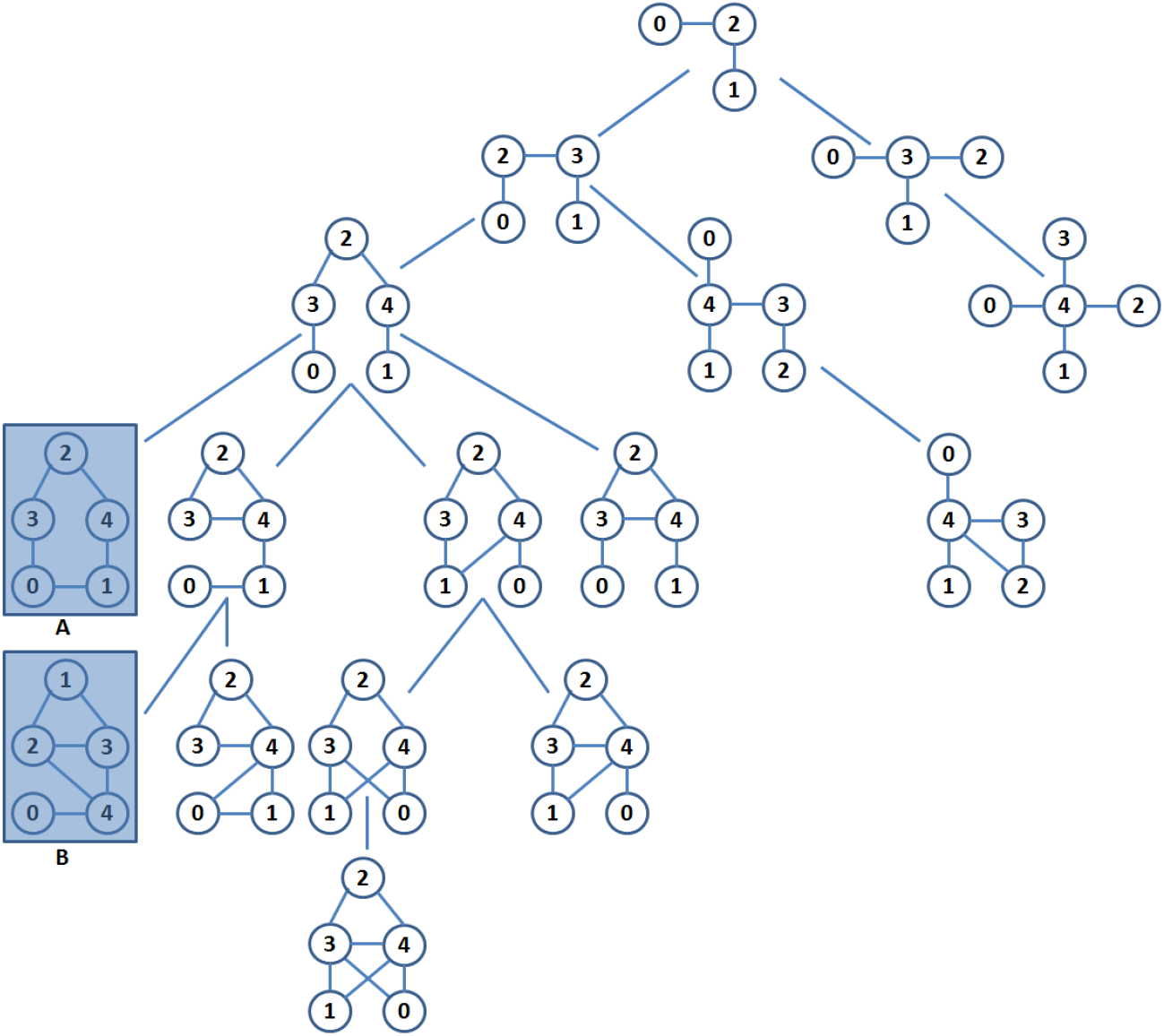
Dynamic expansion tree T_5_ for size-5 motifs

Two shaded node in the dynamic expansion tree (Figure 2) represents subgraphs whose appearances in the target network is less than the frequency threshold. Hence the sub trees rooted with these two nodes are pruned without further expansion.

### 3.3. Vertex Addition Step

During vertex addition, in the adjacency matrix of parent node one extra row and one extra column are appended. Depending on new vertex to be added a row entry and its corresponding column are set as 1. The new tree is taken as a new child node when it is non isomorphic to its sister node (from all parents). The canonical string and the corresponding mapping are stored in the new node.

**Figure 3.**
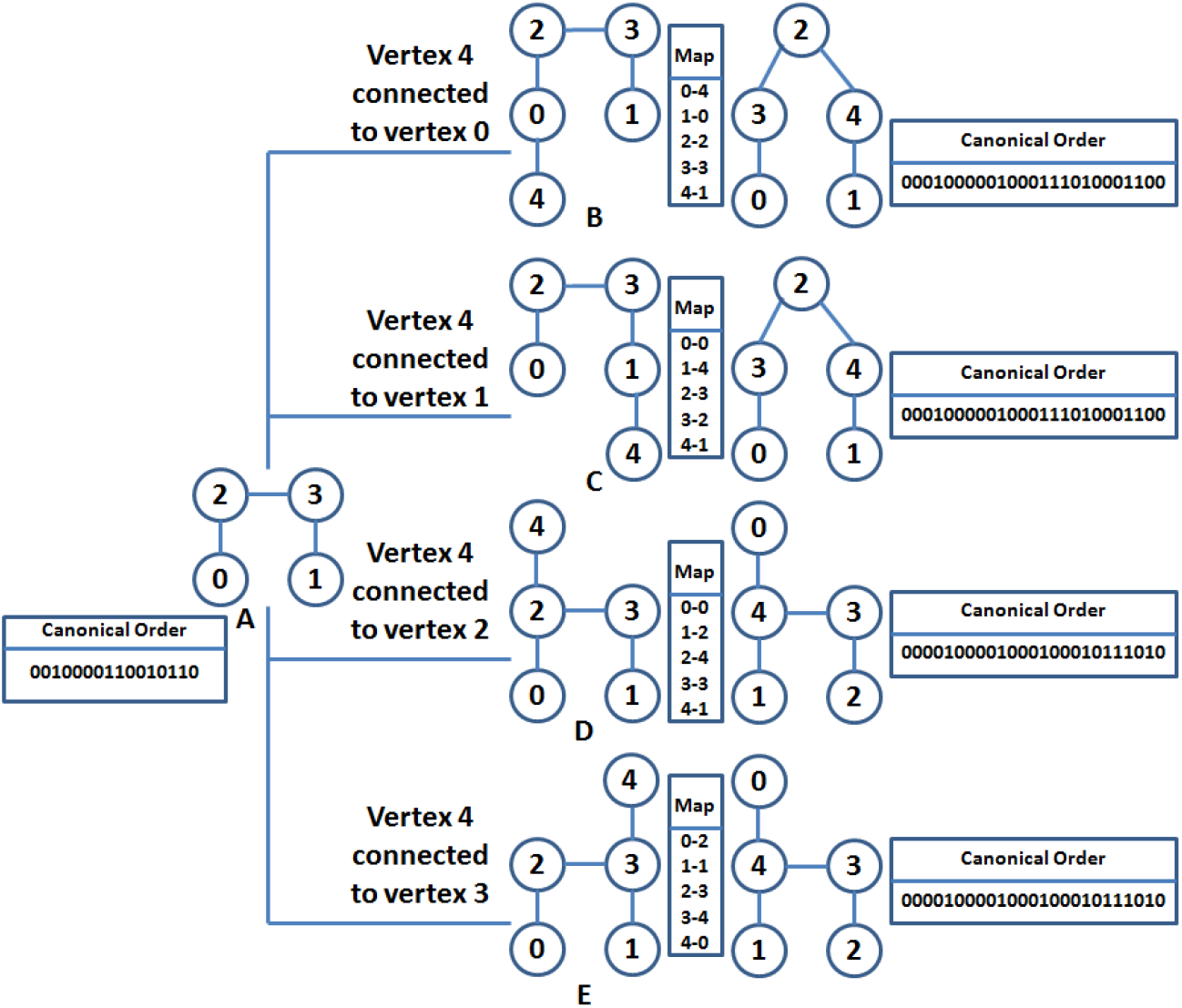
Demonstration of vertex addition step A. Parent graph. B, C, D, E. Child graphs obtained after connecting a new vertex to one of the existing vertex of parent node

In the Figure 3 tree B and tree C results after a new vertex is added to tree A. These are isomorphic with each other as the mapping leads to the same canonical order. Similarly tree D and tree E are also isomer with each other.

### 3.4. Edge Addition Step

Edge addition can be performed by replacing a 0 by 1 in an entry of adjacency matrix of parent node. The new graph is taken as a new child node when it is non isomorphic to its sister node (from all parents). The canonical string and the corresponding mapping are stored in the new node.

**Figure 4.**
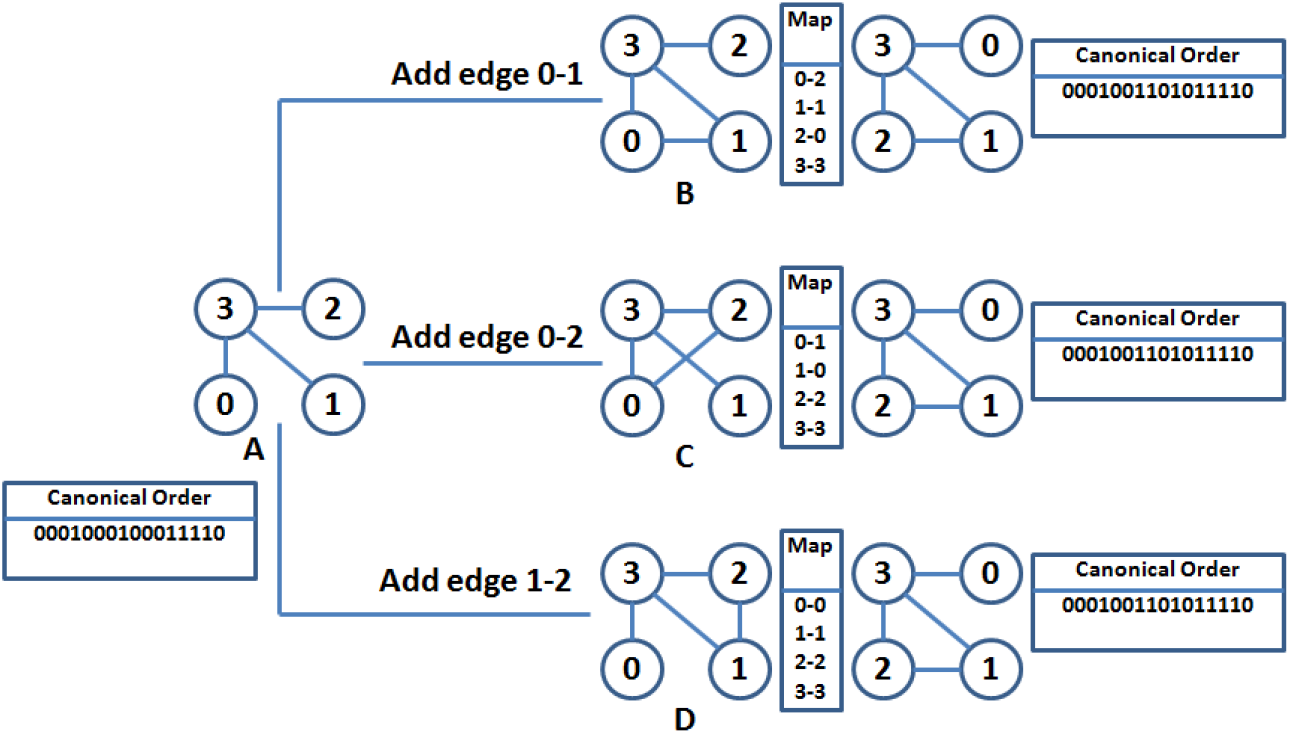
Demonstration of edge addition step A. Parent graph. B, C, D. child graphs obtained after edge addition

Child graphs generated by addition of an edge with the parent are isomorphic with each other (Figure 4) and hence they can be represented by a single node in DET using the mappings shown in Figure 4.

## 4. Proposed Algorithm

In this section, proposed motif discovery algorithm (MDET) is explained. MDET is used to discover statistically significant network motifs in a biological network. The input to the algorithm is a biological network G, a user defined frequency threshold F, a user defined uniqueness threshold Δ, and a user defined maximal network motif size K. The output of the algorithm is a set U of repeated and unique motifs from size 3 to size K.

**Algorithm 1:**
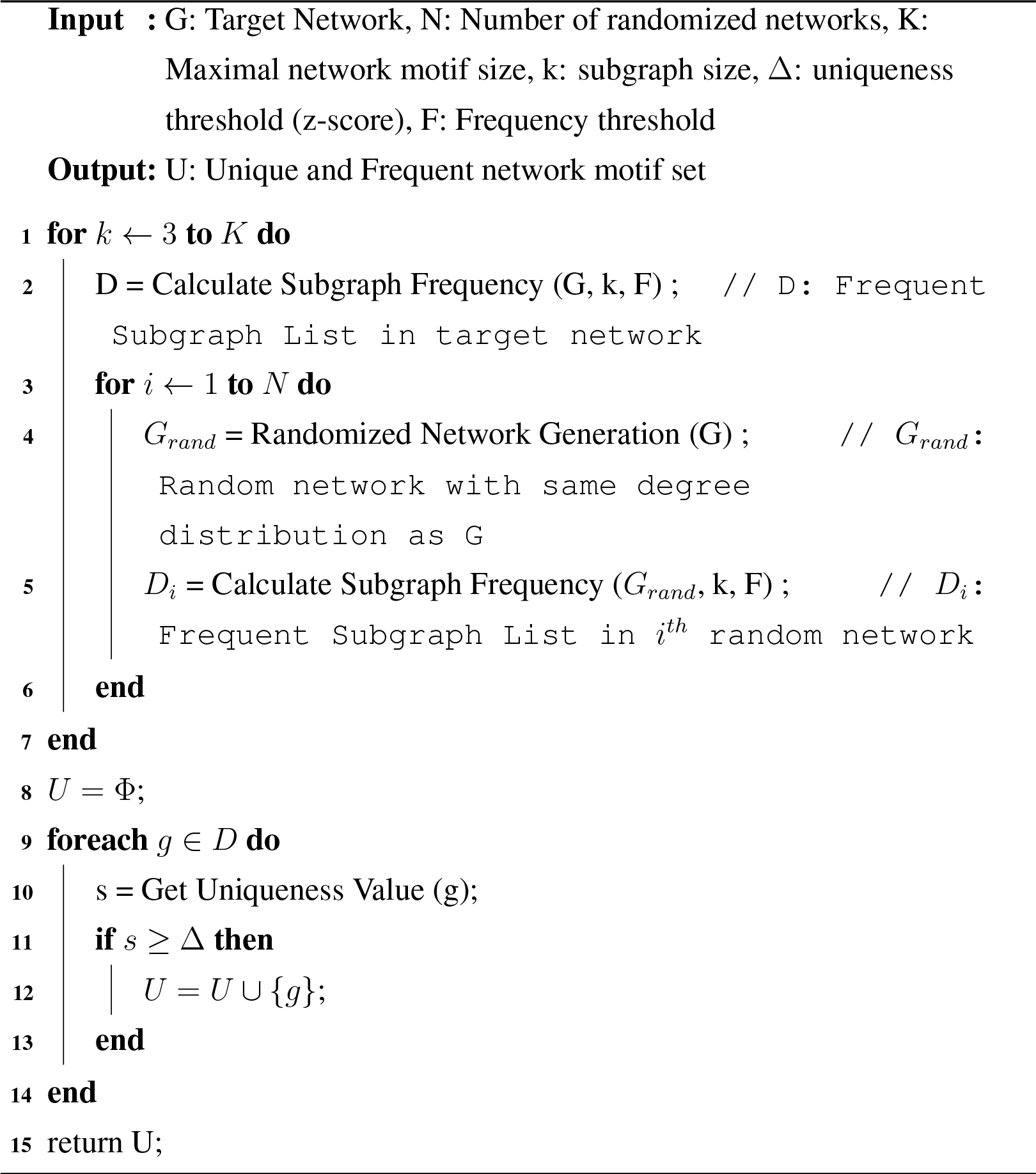
MDET (G, N, K, Δ, F)

The proposed algorithm consists of three main steps. First, we find frequency of repeated subgraphs in the real network (Line 2) using algorithm 2. Then we check the frequency of the repeated subgraphs in the randomized networks (Lines 3-6). Switching method is used to generate random networks [30]. Finally, we determine the unique network motifs from frequent subgraphs (Lines 9-14). We have used z-score of 2 as uniqueness threshold and chosen the frequency threshold as 5% of size of network. Motif-size is taken up to K=15 and statistical significance is measured by taking N=100 random networks. Pseudo-code of the algorithm for discovering network motifs is present in Algorithm 1.

### 4.1. Calculate Subgraph Frequency

Expansion tree is built at the time of frequency calculation. At first, the algorithm construct the root node of the DET and then fetches the size-3 query graph represented by root node of *T_k_* and finds all its embeddings in the target network using algorithm 3. Then it computes the edge disjoint embedding by using Maximum Independent Set (MIS) algorithm and holds these calculated embeddings in memory space for future use. If the frequency of the embeddings exceed frequency threshold then the DET is expanded either by adding vertex or by adding edges depending on required motif size. After that, the query graph at the second level of *T_k_* is fetched and the frequencies of these graphs are calculated either by tree census or by graph census depending on target motif size. Then again edge disjoint embeddings are obtained by Maximum Independent Set (MIS) algorithm and subgraph frequency compared against threshold. This process continues in a depth first order till the pruning criteria is satisfied or a leaf node is obtained where there is no provision for adding new edges. Pseudo-code of the algorithm for calculating frequency of k-size subgraphs is as follows

**Algorithm 2:**
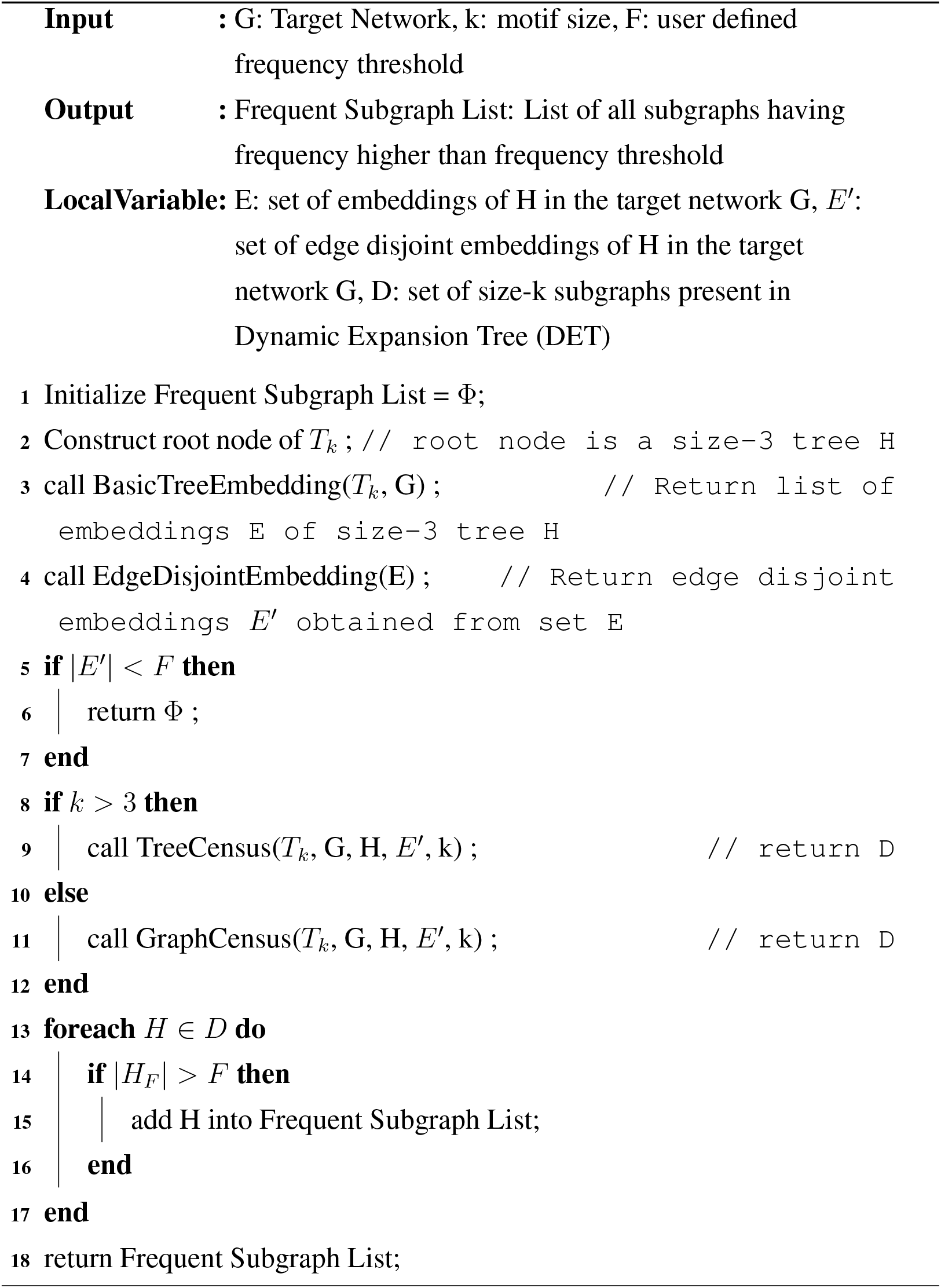
Calculate Subgraph Frequency (G, k, F)

In Algorithm 2, BasicTreeEmbedding function is called in line 3 which returns all the embeddings of size-3 tree. Then in line 4 EdgeDisjointEmbedding function is called which return edge disjoint size-3 tree list using MIS algorithm. Then depending on the input value of k either TreeCensus function or GraphCensus function is called; lines 8-12 perform this task. If the frequency of a size-k subgraph is more than the user defined frequency threshold F then that is added into the Frequent Subgraph List; lines 13-17 perform this task.

### 4.2. Basic Tree Embedding

In this function we are finding all subgraphs isomorphic to the root node of *T_k_*.

**Algorithm 3:**
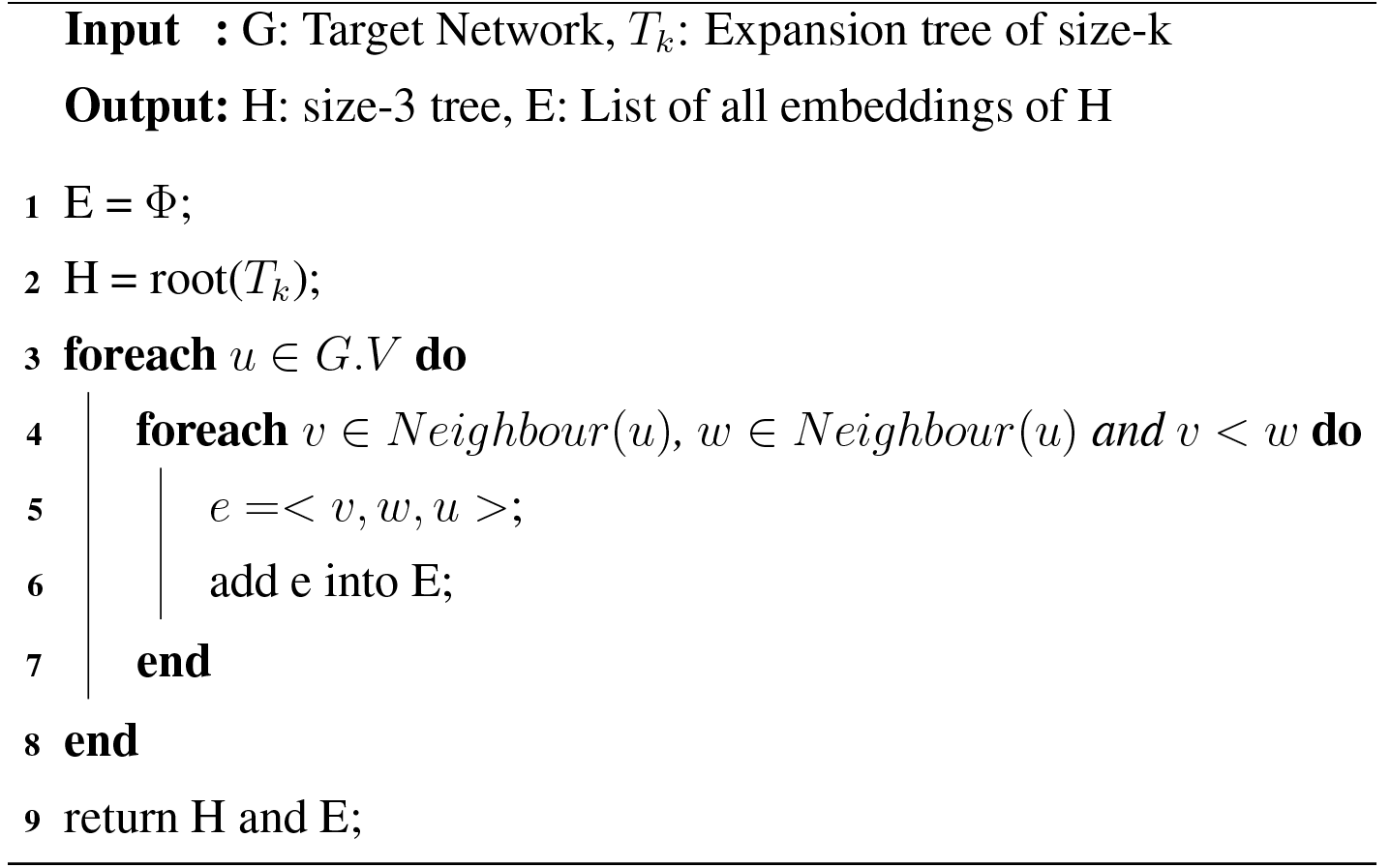
BasicTreeEmbedding(T_*k*_, G)

In Algorithm 3, vertex set of the graph G is denoted as G.V and the neighbouring vertices of a vertex u in the underlying network denoted as Neighbour(u). In line 1 the set of all embeddings of basic tree is initialized to empty set. All the subgraphs both induced as well as none induced of the underlying network are identified and added to set E; lines3-8 perform this task.

### 4.3. Tree Census

This module finds list of all subgraphs isomorphic to the child node using the embeddings of parent node where the child node has one extra vertex and one extra edge than parent node. This procedure can be divided into two phases, i.e. construction phase and expansion phase. In construction phase non isomorphic children are generated from the parent node using vertex addition. In expansion phase frequency of each child is computed and called for expansion if the frequency exceed threshold. Suppose we want to calculate the frequency of a query graph *H’*, we can extract all the embeddings represented by set E corresponding to its parent node H, then enumerate all embeddings in E that can support G and *H’* then store them in *E’*. Let (u, v) be a new edge in *H*^’^ and there exists a vertex f(v) adjacent to f(u) in the target network G, then e can be added to the set *E’*. Where f: *H’* → *G*.

**Algorithm 4:**
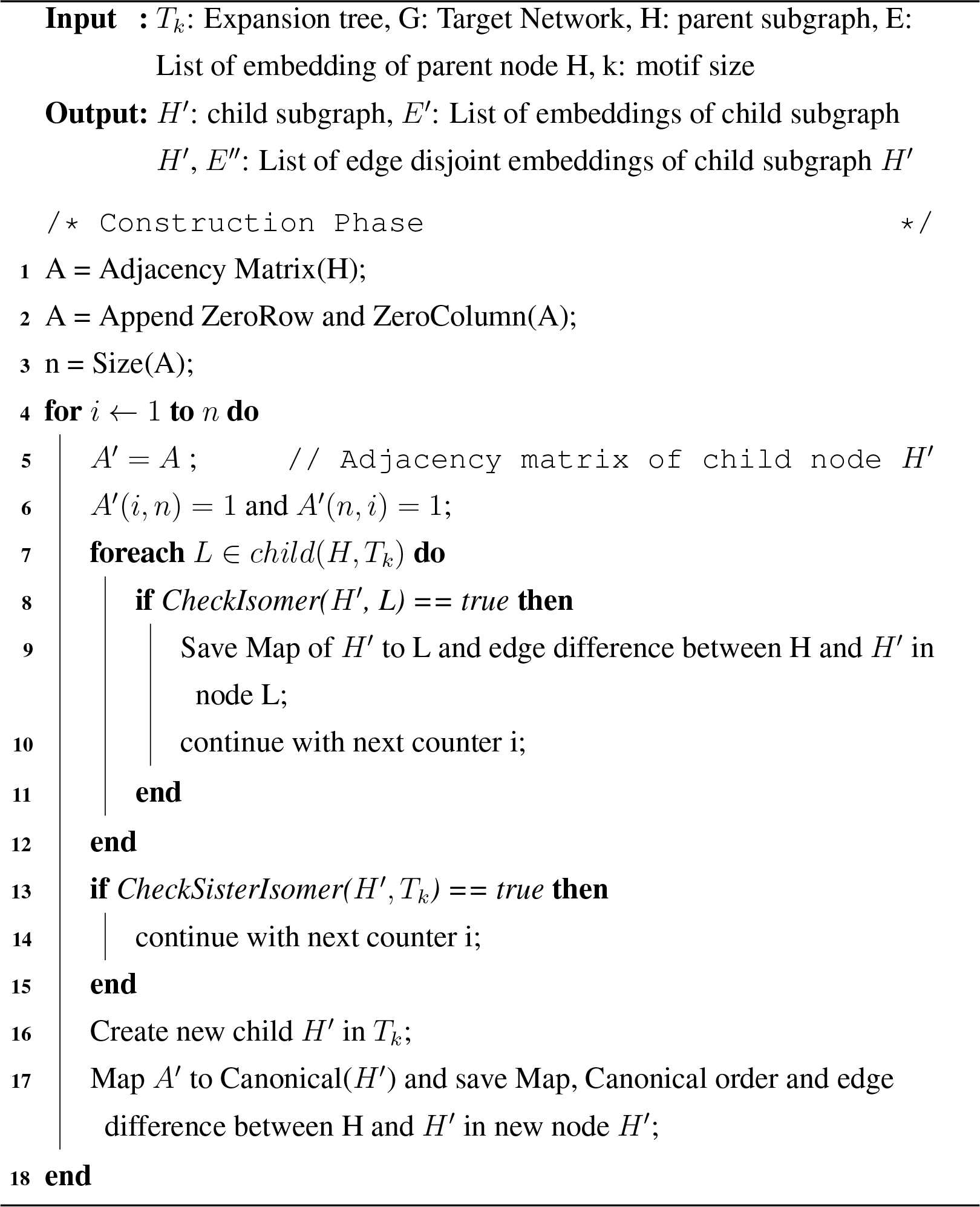
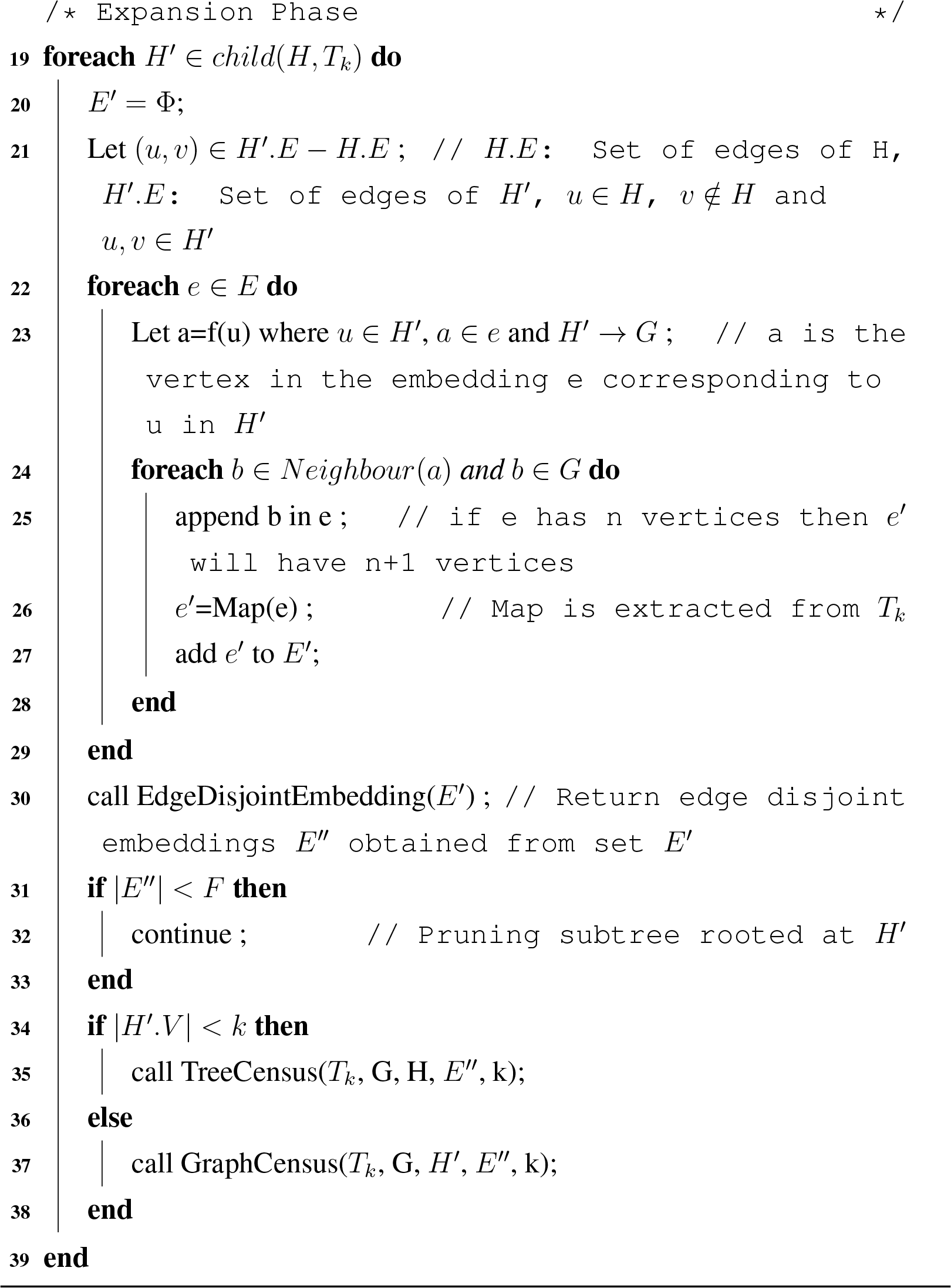
TreeCensus (*T_k_*, G, H, E, k)

Algorithm 4 returns a list of embeddings of child node H in the target network G. Child nodes of expansion tree *T_k_* are created in line 1-18. In line 2 an extra row and an extra column are added with the adjacency matrix of H. In line 6 a new edge is created between an old vertex and the newly added vertex. Lines 7-12 check whether the newly generated graph is an isomer to one of the child created from the same parent, if it is an isomer then the edge difference between parent and new subgraph and the mapping required to convert graph into canonical form are saved in existing child. Then it jumps to the next iteration. Lines 13-15 check whether the newly generated graph is an isomer to any nodes in the expansion tree, if it is an isomer then it jumps to the next iteration. Lines 16-17 creates a new child in the expansion tree corresponding to the new subgraph and store the canonical order of the subgraph along with edge difference between parent and new subgraph and the mapping required to convert the graph into canonical form. Expansion phase starts at line 19. All the child nodes of H present in expansion tree *T_k_* are traversed one by one. The embedding set of child subgraph *H’* is denoted as *E’* and it is initialized to an empty set in line 20. The extra edge need to be added into the parent graph to obtain the child graph is denoted as (u, v). In line 22, the algorithm iterates over all the embeddings of parent graph. Lines 23-28 generates the embeddings of child graph from the embeddings of parent graph. In line 30 edge disjoint embeddings of child graph are obtained from overlap embeddings using MIS algorithm. If the F2 frequency of the child node failed to cross the threshold then the algorithm continues with the next child. This is shown in Lines 31-33. This function recursively call itself until the child graph size reaches to the value k otherwise graph census is called; line 34-38 perform this task.

### 4.4. Graph Census

This module finds list of all subgraphs isomorphic to the child node using the embeddings of parent node, where the child node has an extra edge than the parent node. This procedure can be divided into two phases, i.e. construction phase and expansion phase. In construction phase, non isomorphic children are generated from the parent node using edge addition. In expansion phase, frequency of each child node is computed and called for expansion if the frequency exceeds threshold. Say, we want to calculate the frequency of a query graph *H’*. The embeddings (E) of parent node H extracted first. Then enumerate all embeddings in E that can support G and *H’* and store them in *E’*. Let (u, v) be a new edge in *H’* and there exists an edge (f(u), f(v)) in the target network G, then e can be added to the set *E’*. Where *f*: *H’* → *G*.

**Algorithm 5:**
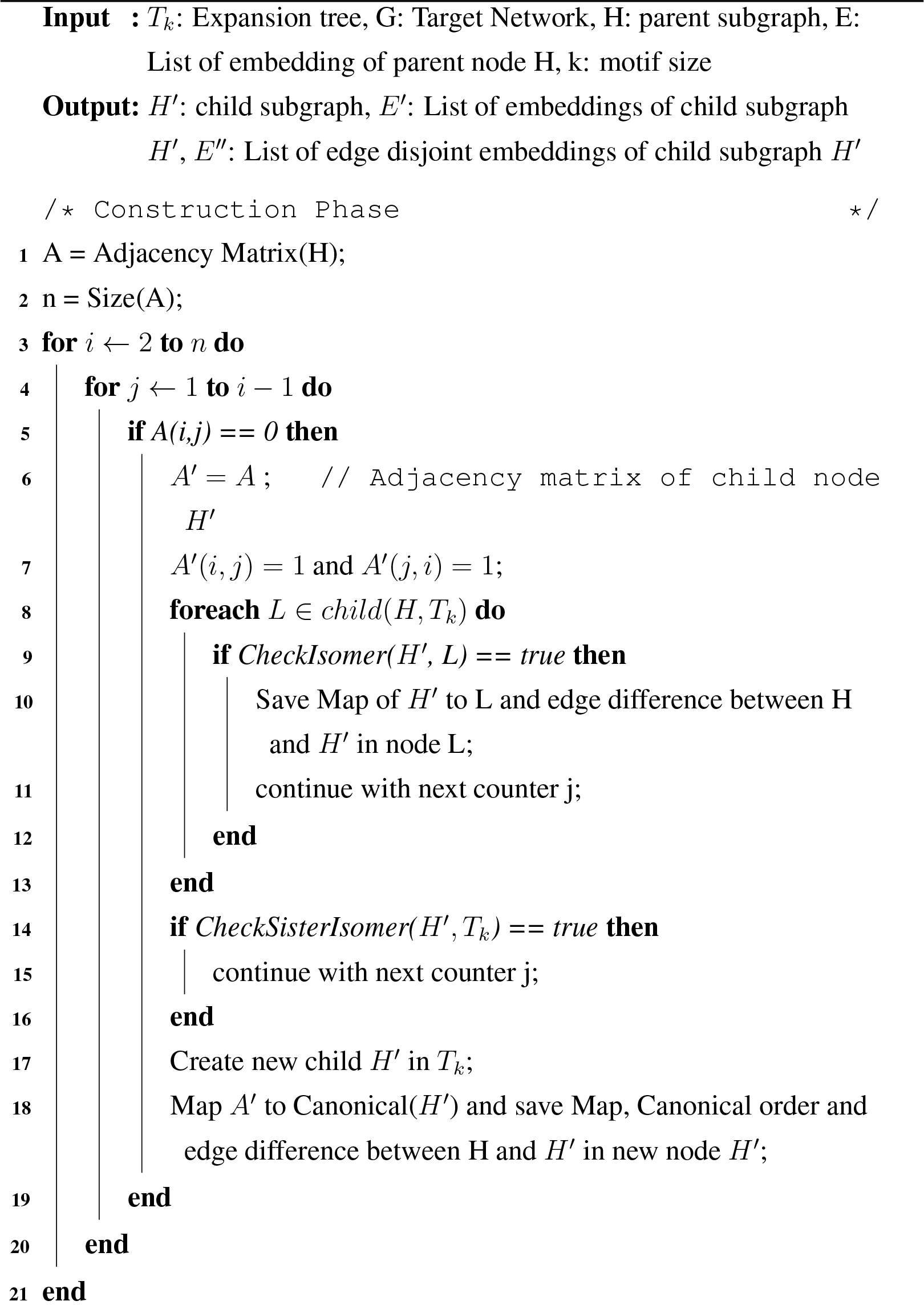
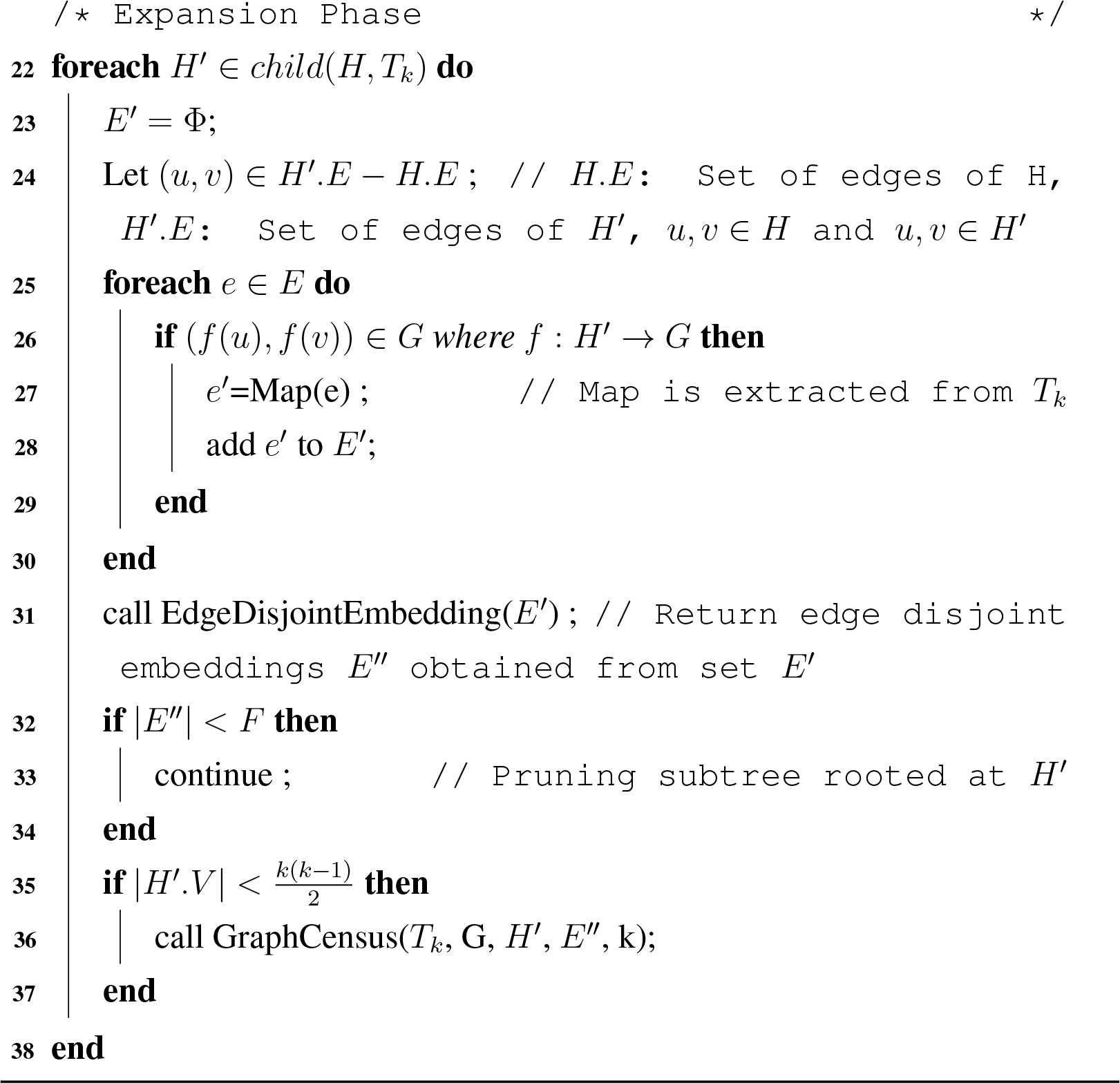
GraphCensus (*T*_k_, G, H, E, k)

Algorithm 5 returns a list of embeddings of child node H in the target network G. Child nodes of expansion tree *T_k_* are created in line 1-21. In line 7 a new edge is created in the adjacency matrix of child node *H’*. Lines 8-13 check whether the newly created graph is an isomer to one of the child created from the same parent, if it is an isomer then the edge difference between parent and new subgraph and the mapping required to convert graph into canonical form are saved in existing child. Then it jumps to the next iteration. Lines 14-16 check whether the newly generated graph is an isomer to any nodes in the expansion tree, if it is an isomer then it jumps to the next iteration. Lines 17-18 creates a new child in the expansion tree corresponding to the new subgraph and store the canonical order of the subgraph along with edge difference between parent and new subgraph and the mapping required to convert the graph into canonical form. Expansion phase starts at line 22. This algorithm iterates over all the child nodes of H present in expansion tree *T_k_*. The embedding set of child subgraph *H’* is denoted as *E’* and it is initialized to an empty set in line 23. The extra edge need to be added into the parent graph to obtain the child graph is denoted as (u, v); line 24 perform this task. In line 25, the algorithm iterates over all the embedding of parent graph. Lines 26-29 perform the task whether the addition of a new edge in the parent embedding support target network or not. In line 27 mapping is done based on the canonical order of the resultant graph after edge addition. In line 31 edge disjoint embeddings of child graph are obtained from overlapped embeddings using MIS algorithm. If the F2 frequency of the child node failed to cross the threshold then the algorithm continues with the next child. This is shown in Lines 32-34. This function recursively call itself until the child graph become a complete graph; line 35-37 perform this task.

### 4.5. Computational Complexity

Here we analyze the time complexity of proposed method. We have expressed the complexity of the algorithms with respect to two parameters n=number of vertices of the target network and k=motif size.

#### 4.5.1. Algorithm 3 (Find all subgraphs isomorphic to size-3 tree)

In this step, embeddings of size-3 tree are generated directly from the degree distribution of nodes. Let *d*(*v_i_*) represent the degree of node *v_i_*. Time complexity of collecting subgraphs isomorphic to size-3 tree is 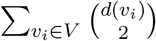. In the worst case *d*(*v_i_*) = *O*(*n*), hence the complexity can be derived as

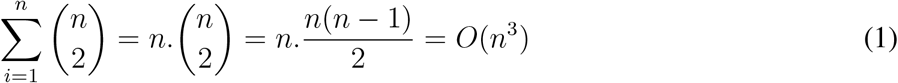

#### 4.5.2. Algorithm 4 (TreeCensus)

In construction phase graph isomorphism check is done which has exponential time complexity. However it is required to check isomorphism only for creating the child nodes. This is limited in numbers and once the child nodes are created no further isomorphism check required in expansion phase. In expansion phase, candidate vertices of parent graph are checked for extension one by one. Let m is the number of candidate vertices for possible extension where m lying between 1 to k. To add a vertex to a candidate vertex all neighbours of the candidate vertex are checked one by one. Thus, the complexity of vertex addition is 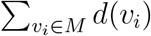. In the worst case *d*(*v_i_*) = *O*(*n*) and the complexity becomes *O*(*nk*) which can be approximately taken as O(n), when *k* << *n*.

#### 4.5.3. Algorithm 5 (GraphCensus)

Similar to TreeCensus here also isomorphism check is required only in construction phase. Hence it is also limited in numbers and does not required in expansion phase. In expansion phase, an edge is added to the parent graph to obtain the child graph. Let m is the number of candidate edges which is lying between 1 to 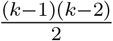.An edge can be added in O(1) time complexity. Thus, the complexity of edge addition is 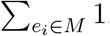. In worst case scenario, the complexity of this algorithm becomes *O*(*k*^2^) which can be approximately taken as O(1).

#### 4.5.4. Algorithm 2 (Calculation of Subgraph Frequency)

Algorithm 3 is called only once. The TreeCensus function is called at max (*k* — 2) times for each embedding of basic tree but most of the embeddings does not appear in child nodes with the increase depth of expansion tree. Similarly the GraphCensus function is called at max 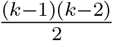 times for each embedding of size-k tree but most of them disappear much before leaf position. In addition to that, the pruning criteria interrupt the growth of most of the branches of expansion tree.

## 5. Results and Discussion

Performance of the proposed motif discovery algorithm is evaluated on real networks taken from MINT database [31]. The run time and number of motifs discovered by proposed algorithm are evaluated across six different networks. In this paper *F*2 measure is used to compute motif frequency and statistical significance of the network motif measured using z-score. Performance of proposed algorithm is compared against MODA.

### 5.1. Data Set and Implementation environment

PPI network of six different organisms from MINT database are used. The details are given in Table 1.

**Table 1.** PPI networks of six different species taken from the MINT database

| Network Name | Network Code | Number of Proteins | Number of Interactions |
| --- | --- | --- | --- |
| Human herpesvirus 8 | hhv8 | 92 | 170 |
| Human herpesvirus 1 | hhv1 | 176 | 353 |
| Escherichia coli | Eco | 402 | 727 |
| Helicobacter pylori | Hpy | 738 | 1643 |
| Rattus norvegicus | Rno | 1825 | 3471 |
| Saccharomyces cerevisiae | Sce | 3187 | 9171 |

Proposed algorithm is implemented in C++ with Intel(R) Xeon(R) E5-2670 Processor 2.3 GHz CPU, 64 GBs of main memory running Redhat Linux operating system.

### 5.2. Run time evaluation

In this section, the running time of proposed motif discovery algorithm is computed on six real PPI networks. The frequency threshold is set as 5% of size of network and z-score is set as 2. F2 measure is used to compute motif frequency. Effect of motif size on the running time is observed by varying motif size from 5 to 15 and the results obtained are shown in Figure 5.

**Figure 5.**
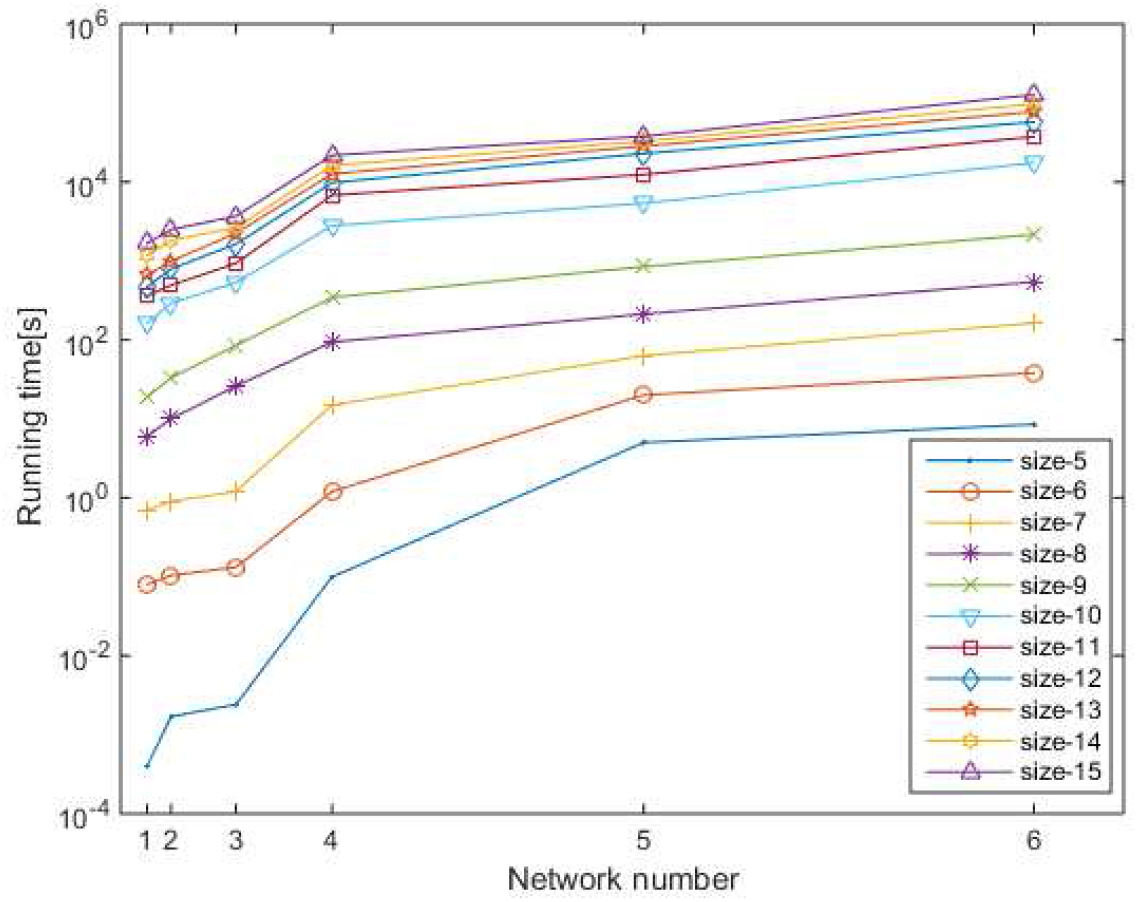
Runtime of MDET for six different PPI networks by varying motif size Network numbers from left to right along x-axis represent hhv-8, hhv-1, eco, hpy, rno and sce respectively. Position of networks along x-axis depends on network size mentioned in Table 1. The running time is measured in seconds.

The behavior of result is a clear indication of scalability of proposed algorithm with respect to graph size and motif size. Proposed algorithm takes only few minutes to run for motif size 5 to 10 even for very large networks and it is limited to few hours for motif size 11 to 15. For higher motif size, run time is more influenced by motif size in compare to size of the network. This behavior is observed due to the number of alternative patterns increases exponentially with respect to motif size. Irrespective of this limitation proposed method is able to discover motif up to size-15 within a practical running time. Table 2 contains the number of motifs found in each of the above network by setting the frequency threshold as 5% of size of network.

**Table 2.** Number of significant motifs for six different PPI networks

| Motif Size | hhv8 | hhv1 | Eco | Hpy | Rno | Sce |
| --- | --- | --- | --- | --- | --- | --- |
| 5 | 10 | 9 | 11 | 9 | 8 | 10 |
| 10 | 5368 | 4219 | 5718 | 3241 | 2816 | 4065 |
| 15 | 8152 | 7529 | 8418 | 6719 | 5245 | 7916 |

### 5.3. Comparison with existing methods

An additional step is introduced in MODA for comparing with proposed algorithm. The experiment has been repeated for three PPI networks and running time is compared between two methods by varying motif size (Figure 6, Figure 7, and Figure 8). MODA and MDET determine the frequency of both induced and non induced subgraphs.

**Figure 6.**
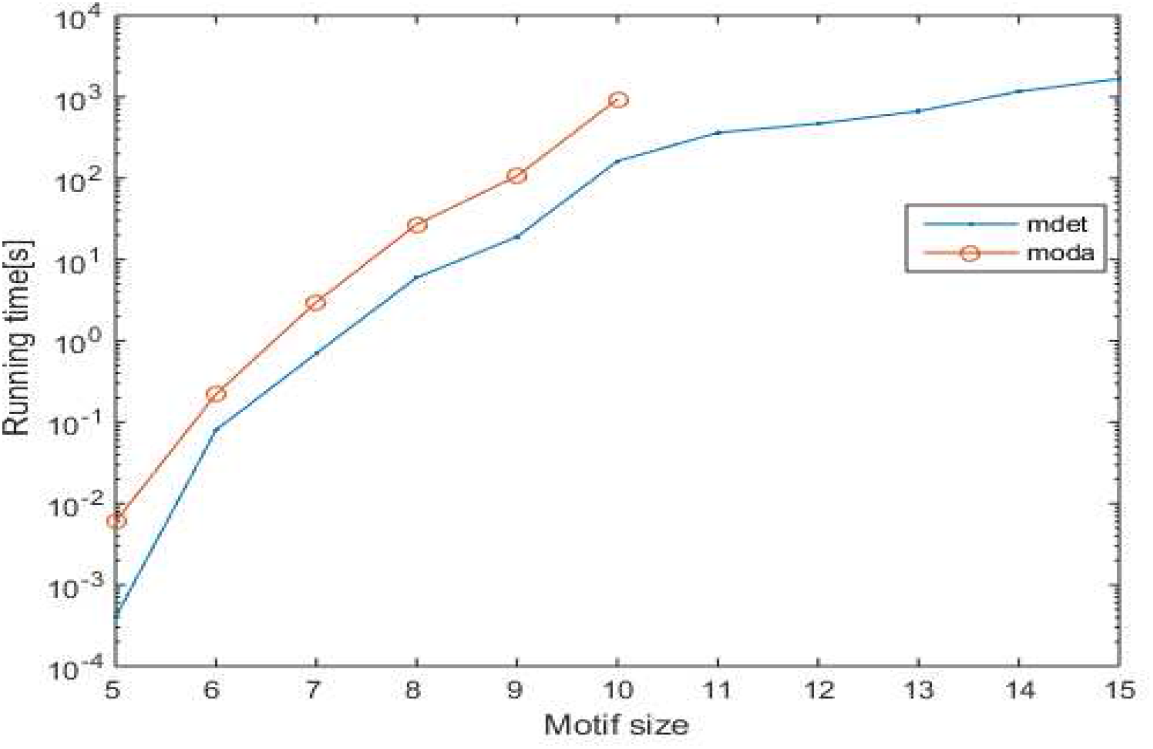
Runtime comparisons between MODA and MDET in Human herpesvirus 8 (hhv8)

**Figure 7.**
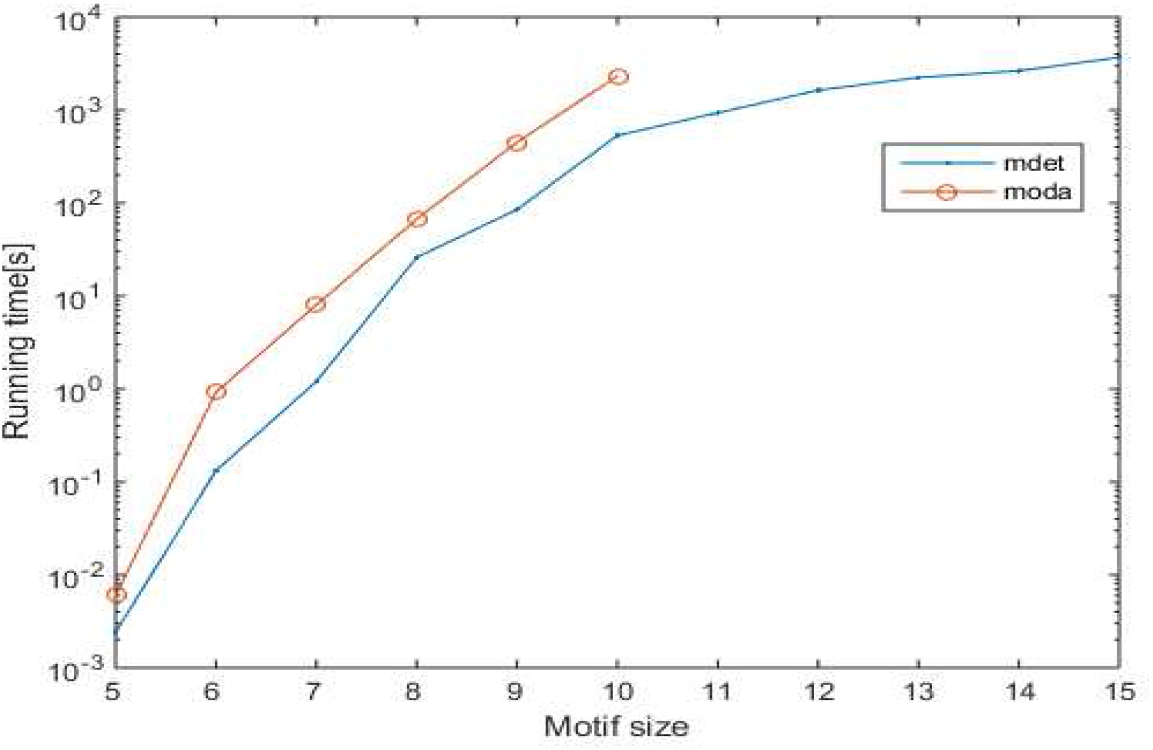
Runtime comparisons between MODA and MDET in Escherichia coli (Eco)

**Figure 8.**
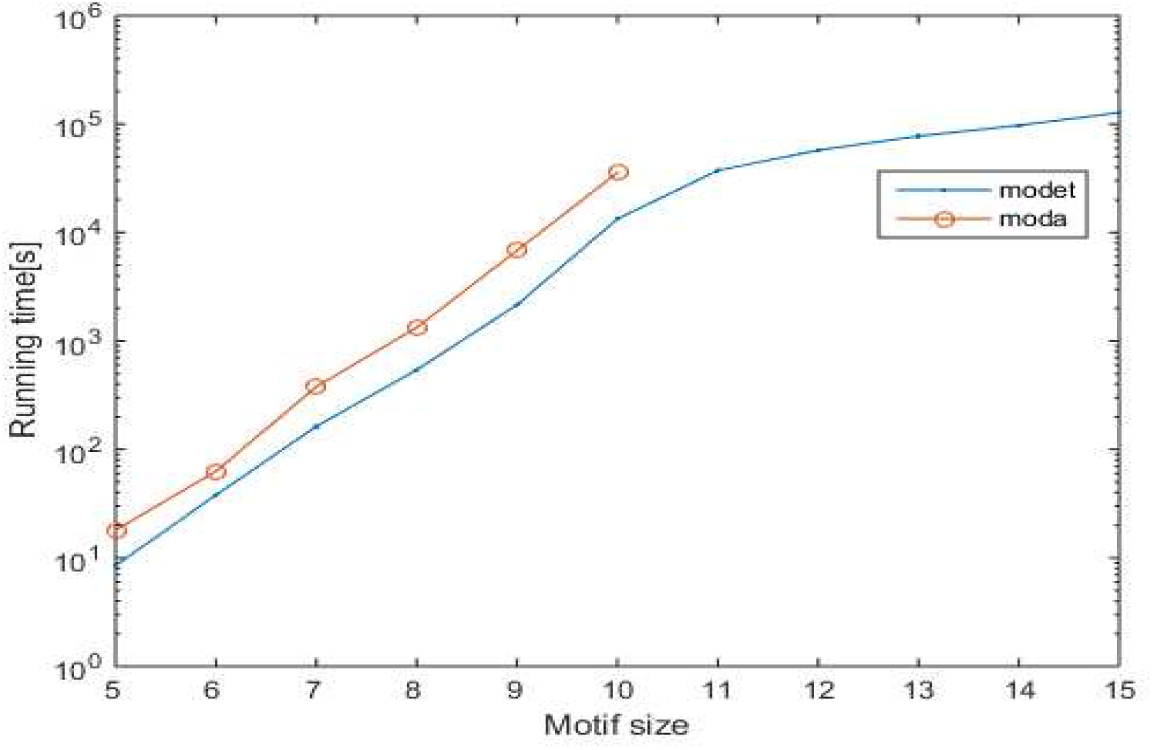
Runtime comparisons between MODA and MDET in Saccharomyces cerevisiae (Sce)

Motif discovery problem exhibit two important characteristics; (1) Number of alternative motif topologies increases exponentially with respect to motif size, (2) the cost of solving subgraph isomorphism also grows exponentially with respect to size of subgraph. Despite these two major concerns, when we increase the motif size the running time of proposed method increases in polynomial order. MODA is able to find motifs up to size-10 only. However proposed algorithm is able to find large motif up to size-15 in practical time bound. MODA uses static expansion tree and hence runs out of space long before it runs out of time. MDET uses dynamic expansion tree, hence this problem is abolished. This fact can be demonstrated with the help of Table 3 and Table 4.

**Table 3.** Number of nodes in static expansion tree Note: Table is prepared using the program geng from McKay’s gtools package [22]

| Motif Size (k) | Cumulative number of non isomorphic trees of size 3 to k | Number of non Isomorphic graphs | Number of nodes in expansion tree |
| --- | --- | --- | --- |
| 3 | 1 | 2 | 2 |
| 4 | 3 | 6 | 7 |
| 5 | 6 | 21 | 24 |
| 6 | 12 | 112 | 118 |
| 7 | 23 | 853 | 865 |
| 8 | 46 | 11117 | 11140 |
| 9 | 93 | 261080 | 261126 |
| 10 | 199 | 11716571 | 11716664 |
| 11 | 434 | 1006700565 | 1006700764 |
| 12 | 985 | 164059830476 | 164059830910 |
| 13 | 2286 | 50335907869219 | 50335907870204 |
| 14 | 5445 | 29003487462848061 | 29003487462850347 |
| 15 | 13186 | 31397381142761241960 | 31397381142761247405 |

Sum of the number of non isomorphic trees starting from size 3 to size k are listed in column 2. This also represents the number of internal nodes in the expansion tree *T*_*k*_. Number of non isomorphic subgraphs of size k are listed in column 3. Total number of nodes in the expansion tree is obtained by adding column 3 with previous row entries of column 2. It can be observed from this table that up to motif size 10, space requirement of the expansion tree is less than 1 GB. But beyond motif size 10 space requirement increases exponentially, and it is impractical to build a static tree for running MODA. However, in a dynamic expansion tree nodes are generated on demand basis. Hence it is quite less than the number of nodes specified in the Table 3.

In Table 4, number of nodes between static and dynamic expansion tree are compared for Escherichia coli network. A uniqueness threshold that is 5% of the size of the network is used. As F2 measure satisfies downward closure property, a node is not further expanded if the frequency of the subgraph less than the threshold value. So most of the branches of DET pruned well before the maximal depth in contrast to static expansion tree. It is observed from Table 4 that number of nodes in static expansion tree increases exponentially with respect to motif size where as in case of dynamic expansion tree it increases linearly. Hence space limitation can be eliminated in MDET method by the use of dynamic expansion tree.

**Table 4.** Comparison between number of nodes in static and dynamic expansion tree Note: Escherichia coli network is used to prepare this table

| Motif Size | 3 | 4 | 5 | 6 | 7 | 8 | 9 | 10 |
| --- | --- | --- | --- | --- | --- | --- | --- | --- |
| Number of nodes in static expansion tree | 2 | 7 | 24 | 118 | 865 | 11140 | 261126 | 11716664 |
| Number of nodes in dynamic expansion tree | 2 | 7 | 15 | 56 | 171 | 645 | 2158 | 7292 |

## 6. Conclusion

In this paper, a Motif discovery algorithm using Dynamic Expansion Tree (MDET) is proposed. The root of the expansion tree is a size-3 tree and the tree is expanded iteratively by addition of graph elements in each successive level. The dynamic expansion tree used in this algorithm is truncated when the frequency of the subgraph fails to cross the predefined threshold. This pruning criterion in DET reduces the space complexity significantly. Pattern growth approach is used in this motif centric algorithm that eliminates costly isomorphism tests. Running time of the proposed algorithm is evaluated by varying motif size and size of the target network. Our implementation results on PPI networks from MINT database indicate that proposed algorithm is significantly faster than the existing motif discovery algorithms and it can able to discover large motifs up to size-15 within few hours. Dynamic expansion tree eliminates memory limitation of static expansion tree. But the space requirement can be further reduced by taking a balanced dynamic expansion tree instead of a simple expansion tree. Motif discovery using balanced dynamic expansion tree can be explained in future.

## Acknowledgements

The authors acknowledge the support by the FIST project of Department of Science and Technology, Govt. of India, the Department of Computer Science and Engineering, IIIT Bhubaneswar, for providing all facilities and guidance.

## 7. References

[1] R. Milo, S. Shen-Orr, S. Itzkovitz, N. Kashtan, D. Chklovskii, U. Alon, Network motifs: Simple building blocks of complex networks, Science 298 (5594)(2002) 824–827.

[2] A. Vazquez, R. Dobrin, D. Sergi, J. Eckmann, Z. Oltvai, A. Barabasi, The topological relationship between the large-scale attributes and local interaction patterns of complex networks, Proc Natl Acad Sci (PNAS) 101 (52)(2004) 17940–17945.

[3] R. Milo, S. Itzkovitz, N. Kashtan, R. Levitt, S. Shen-Orr, I. Ayzenshtat, M. Sheffer, U. Alon, Superfamilies of evolved and designed networks, Science 303 (5663) (2004) 1538–1542.

[4] S. McGee, C. Tibiche, M. Trifiro, E. Wang, Network analysis reveals a signaling regulatory loop in pik3ca-mutated breast cancer predicting survival outcome, Genomics Proteomics & Bioinformatics 15 (2) (2017) 121–129.

[5] R. Gupta, S. Fayaz, S. Singh, Identification of gene network motifs for cancer disease diagnosis, IEEE Distributed Computing, VLSI, Electrical Circuits and Robotics (DISCOVER) (2016) 179–184.

[6] J. Mullen, S. Cockell, H. Tipney, P. Woollard, A. Wipat, Mining integrated semantic networks for drug repositioning opportunities, PeerJ.

[7] L. Li, Z. Wang, P. He, S. Ma, J. Du, R. Jiang, Construction and analysis of functional networks in the gut microbiome of type 2 diabetes patients, Genomics Proteomics & Bioinformatics 14 (5) (2016) 314–324.

[8] F. Schreiber, H. Schwobbermeyer, Frequency concepts and pattern detection for the analysis of motifs in networks, Transactions on Computational Systems Biology III 3737 (2005) 89–104.

[9] M. Kuramochi, G. Karypis, Finding frequent patterns in a large sparse graph, Data Mining and Knowledge Discovery 11 (3) (2005) 243–271.

[10] W. Kim, M. Diko, K. Rawson, Network motif detection: Algorithms, parallel and cloud computing, and related tools, Tsinghua Science and Technology 18 (5) (2013) 469–489.

[11] E. Wong, B. Baur, S. Quader, C. Huang, Biological network motif detection: principles and practice, Briefings in Bioinformatics 13 (2) (2011) 202–215.

[12] M. R. Garey, D. S. Johnson, Computers and Intractability: A Guide to the Theory of NP-Completeness, W. H. Freeman & Co, 1990.

[13] S. A. Cook, The complexity of theorem-proving procedures, Proceedings of the third annual ACM symposium on Theory of computing 18 (1971) 151–158.

[14] N. Tran, S. Mohan, Z. Xu, C. Huang, Current innovations and future challenges of networkmotif detection, Briefings in Bioinformatics 16 (3) (2014) 497–525.

[15] J. Luo, L. Ding, C. Liang, N. Tu, An efficient network motif discovery approach for co-regulatory networks, in IEEE Access 6 (2018) 14151–14158.

[16] G. Ciriello, C. Guerra, A review on models and algorithms for motif discovery in protein-protein interaction networks, Brief Funct Genomic Proteomic 7 (2) (2008) 147–156.

[17] L. Parida, Discovering topological motifs using a compact notation, Journal of Computational Biology 14 (3) (2007) 300–323.

[18] S. Wernicke, F. Rasche, Fanmod: a tool for fast network motif detection, Bioinformatics 22 (9) (2006) 1152–1153.

[19] S. Wernicke, A faster algorithm for detecting network motifs, Algorithms in Bioinformatics 3692 (2005) 165–177.

[20] N. Kashtan, S. Itzkovitz, R. Milo, U. Alon, Topological generalizations of network motifs, Physical Review E (Statistical, Nonlinear, and Soft Matter Physics) 70 (3).

[21] S. Wernicke, Efficient detection of network motifs, IEEE/ACM Transactions on Computational Biology and Bioinformatics 3 (4) (2006) 347–359.

[22] B. McKay, Practical graph isomorphism, Congressus Numerantium 30 (1981) 45–87.

[23] J. Grochow, M. Kellis, Network motif discovery using subgraph enumeration and symmetry-breaking, Research in Computational Molecular Biology 4453 (2007) 92–106.

[24] Z. Kashani, H. Ahrabian, E. Elahi, A. Nowzari-Dalini, E. Ansari, S. Asadi, et al., Kavosh: a new algorithm for finding network motifs, BMC Bioinformatics 10 (1) (2009) 318.

[25] S. Omidi, F. Schreiber, A. Masoudi-Nejad, Moda: An efficient algorithm for network motif discovery in biological networks, Genes & genetic systems 84 (5) (2009) 385–395.

[26] C. Liang, Y. Li, J. Luo, Z. Zhang, A novel motif-discovery algorithm to identify co-regulatory motifs in large transcription factor and microrna co-regulatory networks in human, Bioinformatics 31 (14) (2015) 2348–2355.

[27] R. Elhesha, T. Kahveci, Identification of large disjoint motifs in biological networks, BMC Bioinformatics 17 (2016) 408.

[28] W. Lin, X. Xiao, X. Xie, X. L. Li, Network motif discovery: A gpu approach, IEEE Transactions on Knowledge and Data Engineering 29 (3) (2017)513–528.

[29] Y. Chen, Y. Chen, An efficient sampling algorithm for network motif detection, Journal of Computational and Graphical Statistics.

[30] R. Milo, N. Kashtan, S. Itzkovitz, M. Newman, U. Alon, On the uniform generation of random graphs with prescribed degree sequences, arXiv:cond-mat.stat-mech.

[31] A. Chatr-Aryamontri, A. Ceol, L. Palazzi, G. Nardelli, M. Schneider, L. Castagnoli, G. Cesareni, MINT: the molecular interaction database, Nucleic Acids Research 35 (2007) 572–574.

